# Lipogenic gene expression and substrate sensitivity in the bovine mammary gland shape milk fat composition

**DOI:** 10.64898/2026.08.31.748247

**Authors:** Noam Tzirkel-Hancock, Michal Rakover, Roni Tadmor-Levi, Nurit Argov-Argaman

## Abstract

Milk fat is produced by mammary epithelial cells (MEC) through a conserved mechanism shared among all fat-producing cells across biological kingdoms. Although highly conserved, different tissues and organisms produce distinctive fat compositions. Notably, ruminant milk fat is characterized by enrichment in short and medium chain fatty acids. We hypothesized that this unique profile is driven by MEC-specific metabolic characteristics related to their response to lipogenic substrates. To study this, we compared bovine MEC and udder-derived fibroblasts in terms of their lipogenic capacity and fatty acid composition when exposed to lipogenic building blocks. When exposed to acetate, MEC showed coordinated upregulation of acyl-CoA short-chain synthetase 1 (ACSS1) and diacylglycerol transferase (DGAT), while expression of acyl-CoA synthetase long-chain 1 (ACSL1) decreased. Medium chain fatty acids were also elevated in acetate-treated MEC and not in fibroblasts. The role of ACSS1 in the production of medium chain fatty acids in MEC was confirmed by knockdown experiments. Metabolomics analysis showed that in MEC acetate treatment triggered a broad metabolic response, primarily amino acids catabolism, energy and polar lipid metabolism.

Collectively, these findings demonstrate effective utilization of acetate for *de novo* fatty acid synthesis in MEC with preferred tendency to produce medium chain fatty acids.

## 1. Introduction

In all mammals, milk fat is synthesized by MEC and secreted in a unique structure termed the milk fat globule (MFG). The MFG contains a triglyceride core, which consists of over 95% of total milk fat, enveloped by 3 layers of polar lipids, proteins and carbohydrates derived from the endoplasmic reticulum (ER) and plasma membrane of MEC ^1^. The synthesis of triglycerides starts between the leaflets of the ER, where fatty acids are sequentially esterified to a glycerol backbone. Fatty acids utilized for this lipogenic step are either synthesized *de novo* within the MEC cytoplasm or absorbed from the circulation as preformed fatty acids ^2^. While the *de novo* synthesized fatty acids are typically short and medium chain fatty acids with a carbon chain length of up to 16 atoms, the preformed are long chain fatty acids with a carbon chain length equal to or longer than 16 atoms ^3^.

Bovine milk fatty acids are comprised of approximately 25% short and medium chain fatty acids with a carbon chain length of up to and including 14 atoms. This composition is profoundly different from that of other lipogenic tissues in the same organism, such as the liver and adipose. For example, in the bovine adipose tissue, short chain fatty acids are under detectable levels and the shortest medium chain fatty is myristic acid, making up only 2.5% of total fatty acid composition ^4^. Furthermore, this highly enriched content of short and medium chain fatty acids in ruminant milk is also unique compared to other milk sources. Breast milk for example, contains only ∼11% medium chain fatty acids, with minimal to no presence of short chain fatty acids ^5,6^. The *de novo* fatty acid synthesis pathway is a highly conserved process in various mammals ^2^. However, the underlying mechanism which facilitates the distinguished composition of bovine milk fat is largely unknown.

In all organisms the central enzyme in *de novo* fatty acid synthesis is Fatty Acid Synthase (FASN), a binding enzyme with seven catalytic domains ^2^ which primarily synthesizes palmitate, a 16 carbon saturated fatty acid. The process of fatty acid synthesis by FASN is initiated by acetyl-CoA and involves elongation cycles with malonyl-CoA units ^7^ provided by acetyl-CoA carboxylase (ACC) ^8^. The FASN product- palmitate - is released from the complex by the thioesterase domain of FASN as a free fatty acid ^9^ and can be further elongated or desaturated to increase the variability of fatty acids within the cell. Thereafter, pending activation with coenzyme A (CoA) ^10^, the fatty acid can be used as a substrate for the synthesis of more complex lipids. Fatty acid activation is facilitated by acyl-CoA synthases (ACS), a family of enzymes with varying substrate affinities depending on fatty acid chain length ^11^. After activation, fatty acids enter downstream utilization steps carried out by a series of enzymes, including glycerol-3-phosphate acyltransferase (GPAT), acylglycerol phosphate acyltransferase (AGPAT), and diacylglycerol transferase (DGAT). While AGPAT and GPAT consist of several isoforms in cow tissues, only one DGAT gene has been annotated in cows ^12^. In the mammary gland of dairy cows, short-chain fatty acids, specifically butyric (C4:0), caproic (C6:0), and caprylic acid (C8:0), are predominantly esterified to the sn-3 position of the glycerol backbone^13^ suggesting specific affinity of DGAT to short chain fatty acids - although no such preference has been shown. This stereospecific structure affects the rheological properties of bovine milk fat ^14^, however, the mechanism responsible for this property is yet unknown.

Several mechanisms were suggested over the years to facilitate the biosynthesis and accumulation of short and medium chain fatty acids as triglycerides, similar to those found in cows’ milk: (i) Non FASN thioesterases; these were described in rats ^15^, rabbits ^16^ and even in certain poultry species ^17^. These enzymes can release fatty acids from FASN prior to reaching 16 carbons. However, no external thioesterase was identified in bovine MEC ^12^. Moreover, Hudsen and Knudsen (1980) showed that pharmacological inhibition of thioesterases did not inhibit the ability of bovine FASN to produce short and medium chain fatty acids, suggesting that the release of these fatty acids by FASN is independent of FASN or external thioesterases. (ii) A tissue specific FASN responsible for generating tissue specific fatty acid profiles. It was suggested that structural differences in two catalytic sites of FASN can support the early chain termination of fatty acids by FASN. These differences may include a limited affinity to long chain fatty acids of the ketoacyl synthase domain of FASN ^18^ and an extended affinity of acyl transferase domain of FASN which allows de-loading of short and medium chain fatty acids from this site ^19^. Given that a single FASN isoform is expressed systemically in cows ^20^, if the FASN structure was the main reason for the ability to enrich milk fat with short and medium chain fatty acids, a similar fatty acid profile would likely be observed in all bovine tissues. Therefore, even if these structural changes in FASN contribute to the formation of short and medium chain fatty acids, it cannot be the only mechanism controlling milk fat composition. (iii) Rapid utilization of fatty acids by downstream enzymes by isoforms that may exhibit a higher affinity for shorter chained fatty acids or a higher expression of such enzymes in the mammary gland compared to other tissues. Indeed, specific isoforms like AGPAT1 and AGPAT6 were found to be differentially expressed in the bovine mammary gland compared to other lipogenic tissues ^12,21^. However, it was not demonstrated if they have specific substrate affinity that can contribute to the formation and accumulation of short and medium chain fatty acids in milk triglycerides. (iv) A higher ratio of acetyl-CoA to malonyl-CoA which was shown to enhance the production of short chain fatty acids in rabbits ^22^ and cows ^23^. In ruminants, the primary source for acetyl-CoA is acetate from rumen fermentation of forages ^24^. In accordance, in a mega analysis it was found that ACSS1 which converts acetate to acetyl-CoA, had higher expression in the mammary gland of lactating cows compared with adipose ^12^. However, the involvement of this enzyme in the unique composition of bovine milk fat was not established up to date and no experimental data is available to support the role of ACSS1 in determining the milk fat profile.

Taken together, the mechanism facilitating the production of the unique composition of ruminant milk fat is yet to be determined. In the present study we hypothesized that a combination of metabolic circumstances unique to ruminants, together with the unique response of lipogenic gene expression to metabolic substrates, lead to the distinct composition of fatty acids and triglycerides in bovine milk. We used a comparative approach between different cell types, isolated from the bovine mammary gland and aimed to compare their metabolic and molecular response to lipogenic stimuli.

## 2. Materials and Methods

### 2.1. Cell Culture and Study Design

Three cell lines were used in the present study; two commercial cell lines for bovine MEC (Mac-T, Bet HaEmek, Biological Industries, Bet HaEmek, Israel, and BME-UV1, Biobanking of Veterinary Resources, Italy) and one bovine udder fibroblast cell line, generously donated by Wilk International Biotechnologies LTD (Rehovot, Israel).

The fibroblast cell line was isolated from a bovine mammary gland primary culture generated as previously described ^25^. An immortalization plasmid was constructed and packed by VectorBuilder, containing a puromycin resistance gene coupled with enhanced green fluorescent protein (Puro-EGFP), bovine telomerase reverse transcriptase (Cow-TERT), and SV40 large T antigen (SV40T). The immortalization plasmid was introduced into a bovine primary mammary cell culture containing both MEC and fibroblasts, using Xfect (Takara Bio, California). Following transfection, cells were subjected to puromycin selection for successfully transfected cells, identifiable by EGFP expression. The selected cell population was then stained using anti-cytokeratin 18 (CK18) antibody (Santa Cruz Biotechnology, CA) to identify epithelial cells and separated by FACS (BD FACSMelody Cell Sorter, BD Sciences, NJ). Non epithelial, CK18- negative cells representing the fibroblast population were sorted and seeded as single cells into individual wells. Fibroblast identity was confirmed by morphology and absence of epithelial markers validated by visualization under an Echo inverted fluorescence microscope (Bico, San Diego, CA).

For the experiments, cells were seeded in 10ml plastic dishes with growth medium consisting of DMEM F-12, 10% FBS, 1% L-Glutamine and 1% antibiotics (penicillin and streptomycin mix) all purchased from Biological Industries (Beit Haemek, Israel) and 0.01% cortisol and 0.01% Insulin both purchased from Sigma Aldrich LTD (Rehovot, Israel). Upon reaching confluence, cells were treated with Trypsin–EDTA solution C (Biological Industries, Beit HaEmek, Israel) and re-plated in 30-mm wells at 60,000 cells per ml for Nile Red and DAPI staining, 60-mm petri dishes at 80,000 cells per ml for RNA extraction, or 120-mm petri dishes at 100,000 cells per ml for lipid extraction. All cell types were seeded at the same densities. Once the cells reached confluence again, they were treated with one or more of the following- 40 mM of Acetate, 150 mM of palmitate, 1nM of siRNA (Sigma Aldrich Israel Ltd. Rehovot, Israel) according to the treatments detailed below. All experiments were conducted at least twice, with three biological replicates per treatment (n=3).

### 2.2. Gene Expression Knockdown using siRNA

For transient knockdown of bovine ACSS1, custom endoribonuclease-prepared small interfering RNA (esiRNA) was designed and synthesized by Sigma-Aldritch Ltd (St. Louis, MO, USA) as part of the MISSION® RNAi product line. The esiRNA was based on the bovine ACSS1 mRNA sequence (GenBank accession number NM_174746.2) and was produced by enzymatic digestion of a long double-stranded RNA corresponding to the target region. The reagent was ordered from Sigma Aldrich LTD (Rehovot, Israel). Cells were treated with a 500 ml mix of DMEM F-12 containing 1nM of siRNA and 2μl of transfection reagent for each 60-mm petri dish. For negative control, scrambled non-targeting esiRNA was used with the same conditions. The mix was added to each plate, along with 5ml of fresh prewarmed growth medium per plate. Cells were incubated for 24 hours for optimal results.

### 2.3. Gene expression analysis

After treatment, medium was removed, and the cells were harvested. RNA was extracted using Gene Elute Mammalian Total RNA miniprep kit (Sigma Aldrich, Rehovot, Israel) according to the manufacturer’s instructions. RNA concentration and quality were determined using a nanodrop device (Thermo Scientific, MA) at 260/280 wavelengths. For cDNA synthesis we used qScript cDNA Synthesis Kit (Quanta Biosciences, Beverly MA) according to the manufacturer’s instructions, using 1μg of RNA per sample.

In order to determine expression levels of the selected genes, we designed specific primers for each gene (**Supplementary Table T1**). Gene amplification reactions were preformed using Quantabio PerfeCTa SYBR Green FastMix qPCR kit (Quanta Biosciences, Beverly MA) according to the manufacturer’s instructions. Reaction efficiency and mRNA quantification in the sample was performed using the software LightCycler® 96 (Roche, Basel, Switzerland), and the ΔΔCt method was used to calculate the relative expression of each gene. The expression of the target genes was normalized to two housekeeping genes (UXT, 18S).

### 2.4. Lipid Extraction and fatty acid composition analysis

Cells were harvested and subjected to cold extraction of lipids and methyl ester generation as previously described ^26^. Briefly, chloroform and methanol mixture (2;1 v/v, Bio-Lab Ltd Jerusalem, Israel) was used to extract total lipids, with heptadecanoic acid (C17:0) used as an internal standard (Sigma Aldrich Israel Ltd. Rehovot, Israel) followed by methyl ester generation by methanolic sulfuric acid (95/5, v/v).

Gas chromatographic analysis was conducted using an Agilent gas chromatograph (Agilent Technologies Inc., Wilmington, DE) fitted with a DB-23 fused-silica capillary column (60 m × 0.25 mm i.d., 0.25-μm film thickness; Agilent Technologies Inc.) as previously described ^26^. Compound identification was conducted by comparing retention times with external standards. The separation process was managed by ChemStation software (Rev. B.04.03, Agilent Technologies Inc.). The area under the curve of each compound was normalized to the internal standard and to the total number of cells in the sample, counted using a CytoSmart Cell Counter (Corning, New York, USA).

### 2.5. Cell Staining

Cells were plated on glass cover slips and grown to confluence. Following treatment, the glass covers were washed with phosphate-buffered saline (PBS) (Biological Industries, Beit HaEmek, Israel), fixed with 1ml of 4% paraformaldehyde for 20 minutes at room temperature and washed three more times. The lipid droplets were then stained with 1 ml of Nile Red (Sigma Aldrich LTD, Rehovot, Israel) at the concentration of 1 µg/mL for 15 minutes at room temperature, followed by three more washes with PBS. Lastly, the cell nuclei were stained with 1 ml of DAPI (Sigma Aldrich LTD, Rehovot, Israel) at the concentration of 1 µg/mL for 5 minutes at room temperature and washed three more times. Cover slips were mounted using a fluorescent mounting medium (Dako, Sigma Aldritch Israel Ltd. Rehovot, Israel). Cells were visualized through an Echo inverted fluorescence microscope (Bico, San Diego, CA).

### 2.6. Flow Cytometry

Following treatment, cells were harvested using 0.25% trypsin-EDTA (Sigma Aldrich Israel Ltd. Rehovot, Israel) and collected by centrifugation. Cells were then washed twice with PBS, fixed in 1 ml of 4% paraformaldehyde for 20 minutes at room temperature, and washed again twice with PBS. Lipid droplets were stained using 1 ml of 1 µg/mL of Nile Red for 10 minutes at room temperature in the dark. Cells were washed twice more with PBS and resuspended in PBS for flow cytometric analysis.

Stained cells were transferred to a 96-well plate and analysed using a flow cytometer (Accuri C6, BD Biosciences, New Jersey) equipped with a 488 nm laser. Nile Red fluorescence was measured in the FL2 (585/40 nm) channel, representing lipid content. Total event count was used for cell number quantification and a minimum of 10,000 events per well were collected. Data was analysed using the instrument software (BD Accuri C6 Plus Software).

### 2.7. Metabolomics - Cell Collection and Analysis

At the end of the treatment cells were harvested using 0.25% trypsin-EDTA and collected by centrifugation. The cell pellets were washed three times with PBS to remove residual medium and serum components. Following the final wash, all remaining supernatant was carefully removed, and the cell pellets were snap-frozen in liquid nitrogen and stored at −80°C until metabolite extraction.

### 2.8. Metabolite extraction

Extraction and analysis of polar metabolites and lipids were performed according to previously described methods by Malitsky et al. (2016) and Zheng et al. (2015), with minor modifications ^27,28^. Cell pellets were extracted with 1 mL of a pre-cooled (−20°C) homogeneous mixture of methanol: methyl-tert-butyl ether (MTBE) at a 1:3 (v/v) ratio. Samples were briefly vortexed and then sonicated for 30 minutes in an ice-cold sonication bath, with brief vortexing every 10 minutes. Subsequently, 0.5 mL of UPLC-grade water (DDW): methanol (3:1, v/v) containing C13- and N15-labeled amino acid standards (Sigma, 767964;(1:1500)) was added. After vortexing thoroughly and centrifuging, the organic phase was discarded. The remaining polar phase underwent a second extraction with an additional 0.5 mL of MTBE. The polar phase was transferred to a separate tube and was dried under a gentle nitrogen stream and stored at −80°C.

### 2.9. LC-MS Analysis of Polar Metabolites

Dried polar extracts were resuspended in 120 µL methanol: DDW (50:50), centrifuged twice, and 70 µL was transferred to HPLC vials. Metabolite profiling was performed using an Acquity I-class UPLC system (Waters) coupled to a Q Exactive Plus Orbitrap™ mass spectrometer (Thermo Fisher Scientific), operating in negative ionization mode. Chromatographic separation was conducted using a SeQuant Zic- pHilic column (150 mm × 2.1 mm) with a SeQuant guard column (20 mm × 2.1 mm) (Merck). Mobile phase B consisted of acetonitrile, while mobile phase A comprised 20 mM ammonium carbonate with 0.1% ammonium hydroxide in DDW: acetonitrile (80:20, v/v). The flow rate was 200 µL/min with the column temperature maintained at 45°C. The gradient conditions were: 0–2 min at 75% B; linear decrease to 25% B by 14 min, maintained until 18 min; increased to 75% B at 19 min, held for 4 min, and maintained at 75% B until 23 min. Injection volume: 2 µL. Mass spectral data were acquired within an m/z range of 70– 1050 using heated electrospray ionization (HESI) in negative mode. Ion source parameters included capillary temperature at 325°C, spray voltage at 3.25 kV, sheath gas flow rate at 40, auxiliary gas flow rate at 10 (arbitrary units), and auxiliary gas temperature at 50°C. MS1 spectra were acquired at a resolution of 35,000 FWHM, with data-dependent MS/MS acquisition conducted using an isolation window of 3 m/z and a resolution of 17,500 FWHM.

### 2.10. Polar metabolites data analysis

Data processing was done using TraceFinder (Thermo Fisher Scientific) software, when detected compounds were identified by retention time, and fragments were verified using an in-house-generated mass spectra library. Relative levels of polar metabolites were normalized to the internal standards and total peak value of the original samples.

Following normalization, significantly altered metabolites were determined between control and acetate-treated MacT cells, based on a p-value threshold of 0.05. Additionally, metabolite heat map and pathway enrichment analysis was performed in MetaboAnalyst 6.0 (Xia Lab, McGill University; www.metaboanalyst.ca) using metabolites that passed the significance threshold (*p ≤ 0.05*). Pathway impact scores were calculated based on topology analysis, and directionality (increase or decrease) was assigned based on the majority trend of metabolites within each pathway. Specific metabolites in the enriched pathways were identified and quantified using MetaCyc database (Bioinformatics Research Group, SRI International, CA, USA).

### 2.11. Statistical Analysis

Statistical analysis was conducted using JMP software version 18.0 (SAS Institute, Cary, NC). Gene expression analyses were performed using one-way ANOVA followed by Tukey–Kramer HSD. Fatty acid composition and metabolite level comparisons were analysed using Student’s t-test. Significant probability was set to 0.05, and tendencies were reported at 0.05 < P ≤ 0.1. Results are reported as mean ± standard error mean. Principal component analysis (PCA) was performed on correlation matrices using JMP to explore patterns in the data and reduce complexity. Data were first normalized, and the main components explaining the variation between samples were plotted to identify group clustering and variable contribution.

## 3. Results

### 3.1. Basal gene expression and fatty acid composition; comparing three cell lines

In order to understand the basal lipogenic capacity of each cell type, gene expression of lipogenic genes was determined in bovine fibroblasts as well as two bovine MEC lines (MacT, BME-UV1). Levels of mRNA were measured for fifteen genes encoding enzymes involved in fatty acids and triglyceride synthesis and compared between cell lines. The list of genes was compiled based on previous research by Tadmor-Levi and Argov-Argaman ^12^, that provided a mega- analysis of gene expression in different lipogenic bovine tissues. Fatty acid composition of each untreated cell line after growing to confluence was also compared (**Figure 1**).

**Figure 1.**
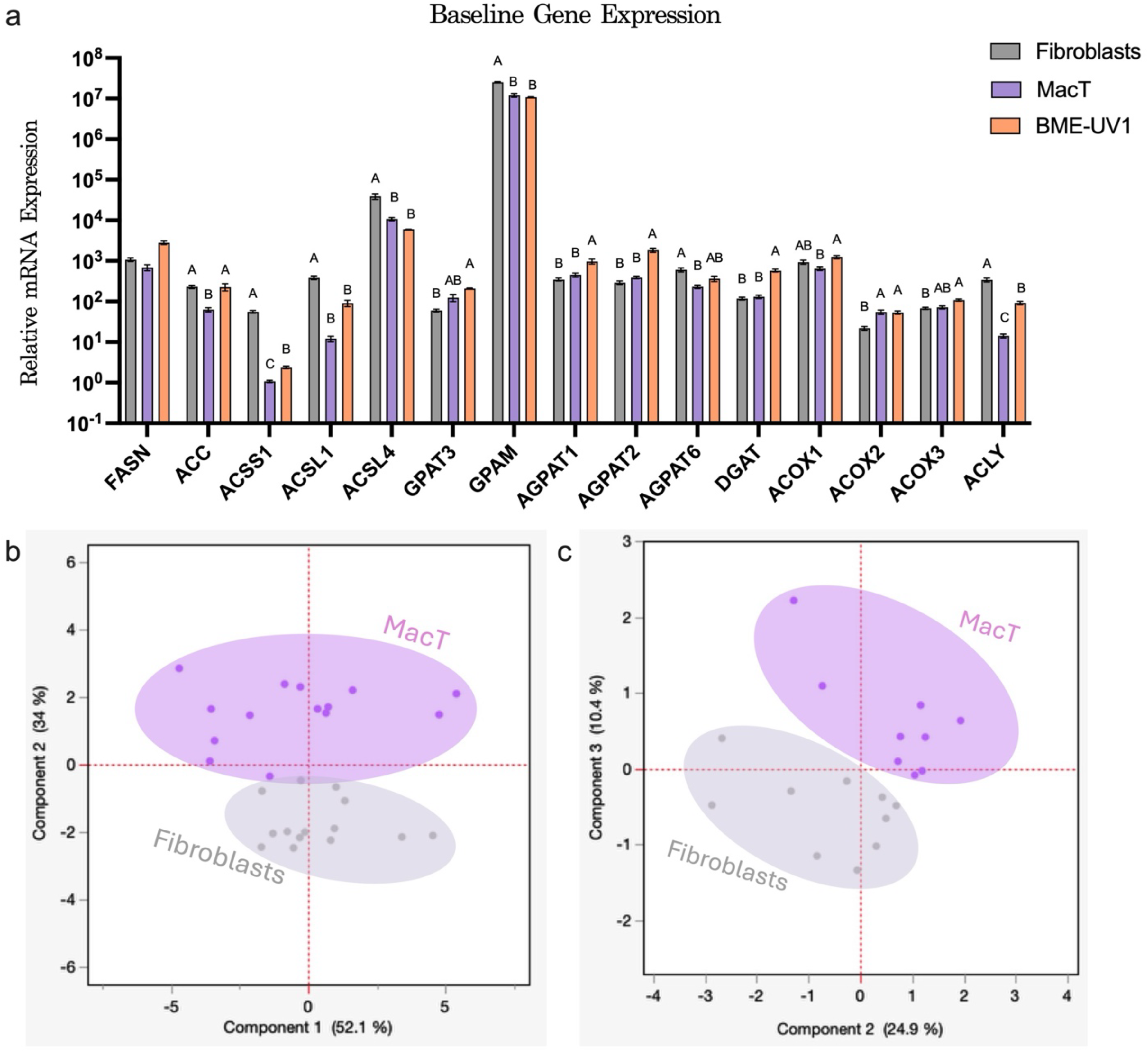
Baseline lipogenic profiles of bovine fibroblasts and MEC. (a) Baseline mRNA expression levels of lipogenic genes in fibroblasts, MacT, and BME-UV1. Expression values are normalized relative to the gene with the lowest expression across all genes and cell types (i.e. ACSS1 in MacT) allowing comparison between cells as well as between genes within the same cell line. Data is presented as mean ± SEM. Different letters indicate significant differences between cell types for each gene (*p* ≤ 0.05). (b) PCA of relative mRNA expression of lipogenesis-related genes in fibroblasts and MacT cells. (c) PCA of cellular fatty acid composition (mol%) in the same cell types.

Across nearly all genes, fibroblasts exhibited a distinct expression profile from both mammary-derived cell types, emphasizing their differences (**Figure 1a**). Notably, genes involved in fatty acid activation (ACSS1, ACSL1, ACSL4) and initial triglyceride synthesis (Glycerol-3-Phosphate Acyltransferase 1, GPAM) were expressed at significantly higher levels in fibroblasts compared to both MacT and BME-UV1, with GPAM being the most significantly expressed gene across all cell lines. In contrast, genes central to *de novo* fatty acid synthesis, such as ACC and ACLY, were more highly expressed in BME-UV1 than in fibroblasts. Additionally, peroxisomal oxidation genes (Peroxisomal acyl-coenzyme A oxidase, ACOX1–3) demonstrated varying expression patterns, with BME-UV1 and MacT expressing higher levels of ACOX2 and ACOX3 than fibroblasts. These differences emphasize that fibroblasts and MEC possess fundamentally distinct expression properties under basal conditions.

PCA of gene expression data revealed clear separation between MacT cells and fibroblasts, with the first principal component explaining 52.1% of the total variance and the second component explaining 34.9% of the variance (**Figure 1b**), highlighting inherent transcriptional differences between these cell types. PCA of fatty acid composition also showed separation between cell types, although explained a lower percentage of the variance, with the second and third principal components explaining 24.9% and 10.4% of the variance, respectively (**Figure 1c**), indicating that lipid profiles reflect and complement the underlying gene expression differences.

### 3.2. MEC and Fibroblast Response to Acetate and Palmitate; Lipogenic Gene Expression

To investigate how fibroblasts and MEC respond to lipogenic substrates we focused on two cell lines, bovine udder fibroblasts and MacT as a representative cell line of MEC. Both cell types were treated with palmitate, acetate, or their combination for 5 or 24 hours and expression of key lipogenic genes was analyzed. Hierarchical clustering of expression data (**Figure 2a**) revealed distinct grouping of fibroblast and MEC samples, indicating that the two cell types have fundamentally different responses to the same metabolic stimuli. ACSS1 and DGAT clustered closely together, particularly in MEC, suggesting coordinated regulation under palmitic acid and acetate exposure, a co-regulation less prominent in fibroblasts.

**Figure 2.**
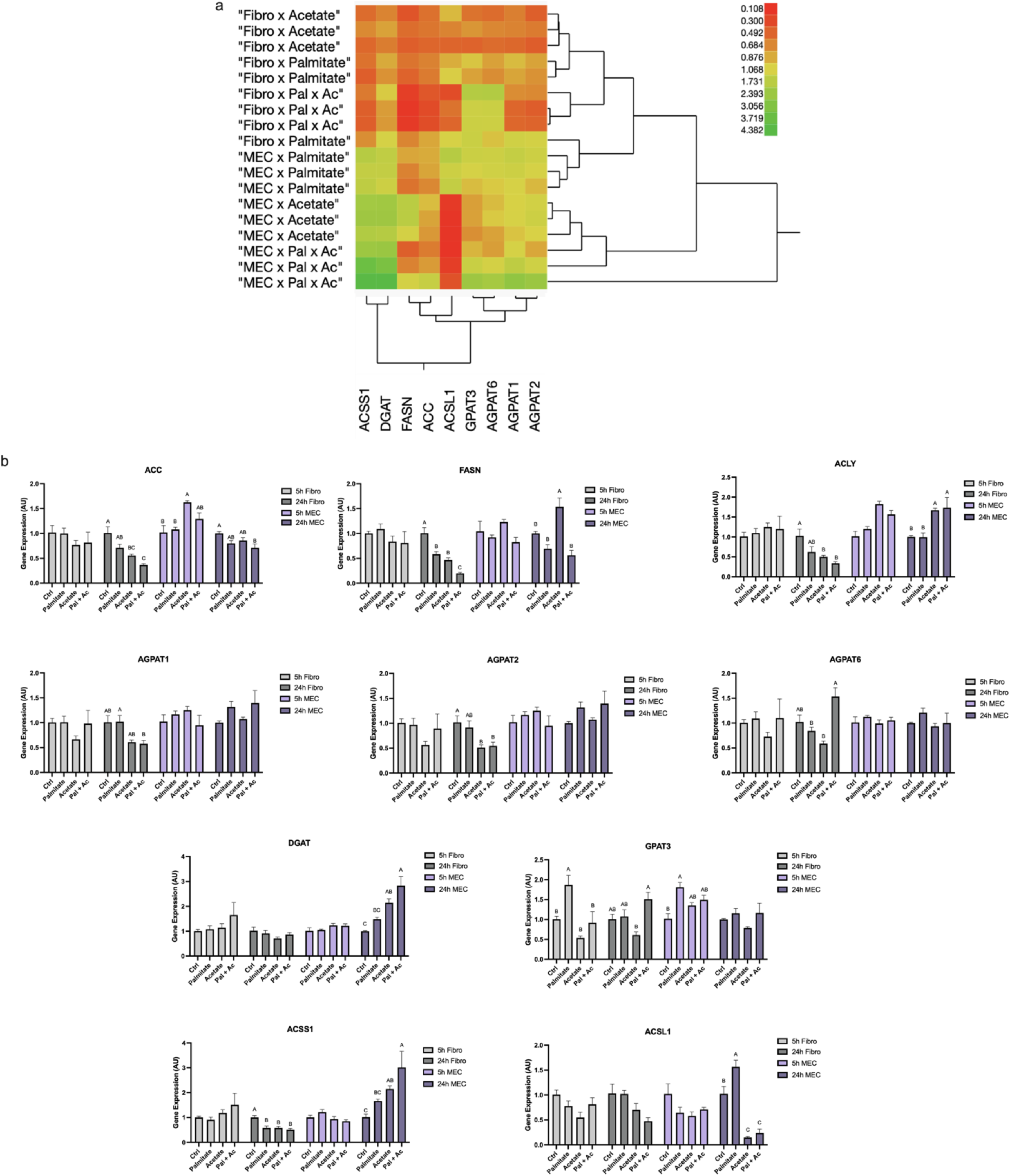
Gene expression responses to acetate and palmitate in bovine fibroblasts and MEC. (a) Hierarchical clustering heat map of lipogenic genes following 24-hour treatment with 150 μM of palmitate, 40 mM of acetate, or both (Pal + Ac) in fibroblasts and MEC. Colour scale represents normalized expression values- red indicates low expression and green indicates high expression, relative to control. (b) Relative bar graphs showing relative mRNA levels of selected genes under each treatment condition (acetate, palmitate or both) and time point (5 or 24 hours). Data represent mean ± SEM. Significant differences (*p* ≤0.05) are indicated by different letters within each gene panel.

The direction of the ACSS1 response to acetate diverged between the two cell types - acetate reduced ACSS1 expression in fibroblasts but increased it in MEC, especially at the 24-hour time point (**Figure 2b**). This finding suggests cell specific acetate utilization. Additionally, ACSL1, an enzyme that activates long- chain fatty acids, was strongly downregulated in MEC upon acetate treatment but remained relatively stable in fibroblasts. DGAT expression was greatly increased in MEC in response to palmitate and acetate, with no significant response in fibroblasts. FASN also responded uniquely in MEC, with a significant increase in response to acetate, while in fibroblasts the expression decreased.

### 3.3. Response to Acetate and Palmitate; Fat content and Fatty Acid Composition

We further determined how fibroblast and MEC responded to treatment with acetate and palmitate in terms of fat content and fatty acid composition (**Figure 3**). In MEC, acetate supplementation increased medium chain fatty acid concentrations, including C12:0 and C14:0, as well as palmitate (C16:0). These changes were not observed in fibroblasts, which exhibited a reduction in C16:0, C18:0 and C18:1n9 cellular content. In contrast, fibroblasts responded strongly to palmitate with an increase of C16:0, an effect not seen in MEC, which maintained a stable fatty acid composition under palmitate stimulus.

**Figure 3.**
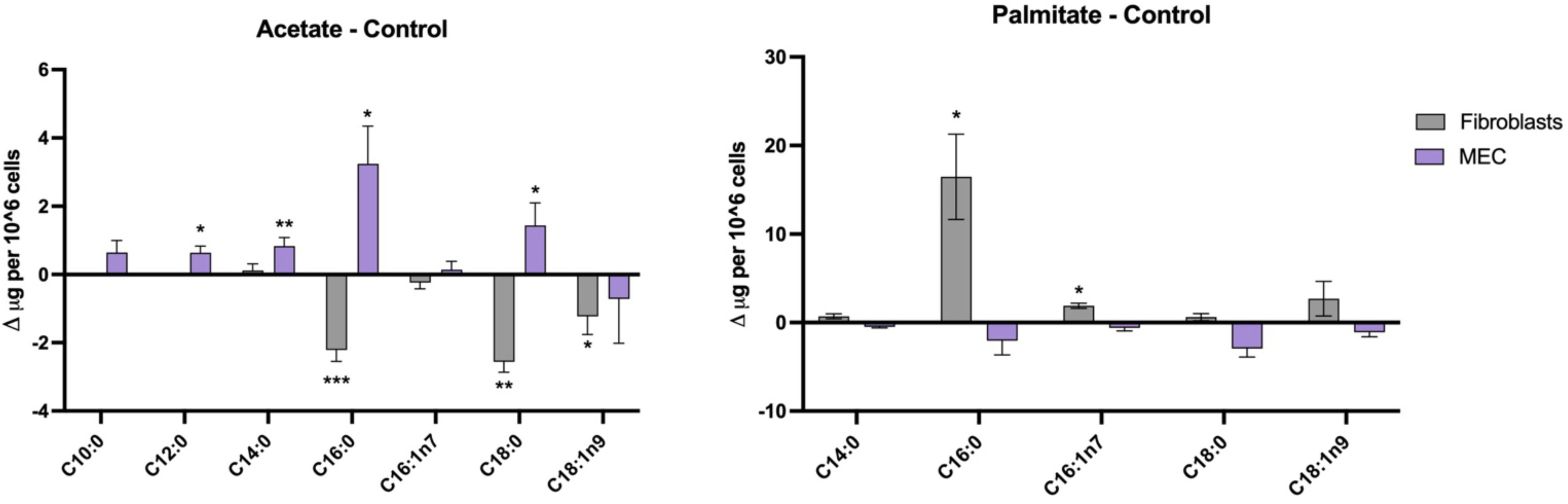
Fatty acid content quantification in fibroblasts and MEC under control, acetate, and palmitate treatment conditions. Total fatty acid content in µg per 1 million cells, in fibroblasts and MEC exposed to 40 mM acetate or 150 mM of palmitate. Values represent the difference (Δ) between acetate or palmitate treated cells and the control groups. Significant differences (p ≤ 0.05) between control and acetate are marked with asterisks.

Intracellular triglyceride content in fibroblasts and MEC was estimated by the intensity of hydrophobic dye (Nile Red) incorporated into lipid droplets. Acetate treatment increased lipid droplet accumulation in MEC, (**Figure 4a**) and induced a clear shift in fluorescence intensity (**Figure 4c**), indicating a significant increase in intracellular lipid content. No shift was observed in fibroblasts (**Figure 4b**), where fluorescence profiles remained comparable between control and acetate treated cells.

**Figure 4.**
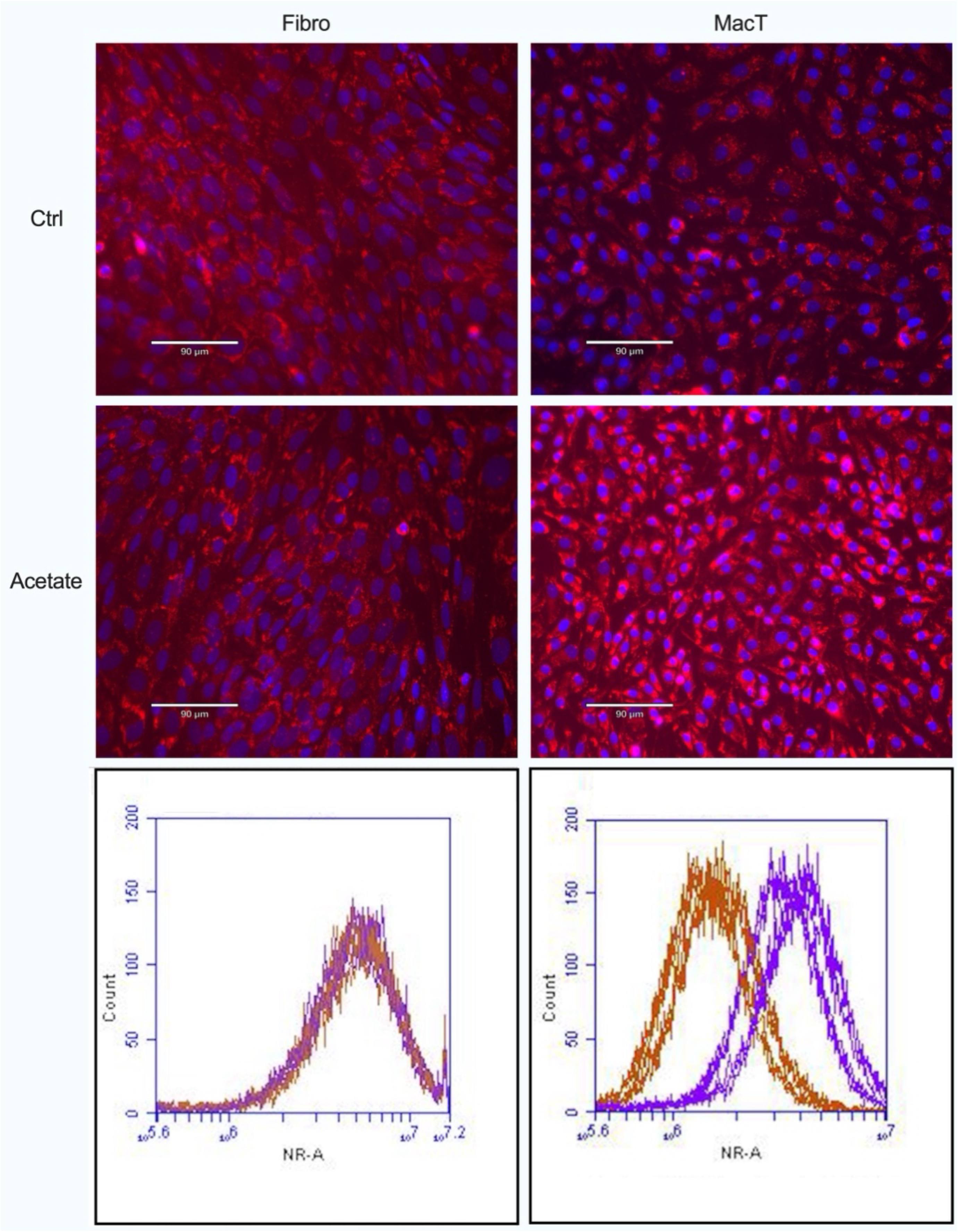
Fluorescence microscopy and flow cytometry analysis of lipid content in fibroblasts and MEC treated with acetate. (a) Representative fluorescence microscopy images of fibroblasts and MEC stained with Nile Red (lipid droplets, red) and DAPI (nuclei, blue), comparing control and acetate (40 mM) treated groups. (b and c) Flow cytometry analysis of Nile Red fluorescence in fibroblasts and MEC, respectively. X-axis: FL2 channel representing Nile Red intensity (lipid content); Y-axis: event count. Purple represents acetate treated cells; brown represents the control group (2 biological repetitions, in triplicates).

### 3.4 ACSS1 role in Fatty Acid Composition Response of MEC

To assess the role of ACSS1 in acetate driven lipid metabolism, we preformed transient knockdown of the ACSS1 gene with esiRNA and subsequent acetate treatment in MEC and fibroblasts. mRNA expression was measured compared to the control group which was treated under the same conditions with scrambled non-targeting esiRNA. ACSS1 expression was significantly reduced following esiRNA treatment, in both MEC and fibroblasts. However, in MEC acetate treatment significantly upregulated ACSS1 expression, while fibroblasts showed no such response (**Figure 5a**). These results were consistent with the response of both cell lines to acetate treatment (**Figure 2b**).

**Figure 5.**
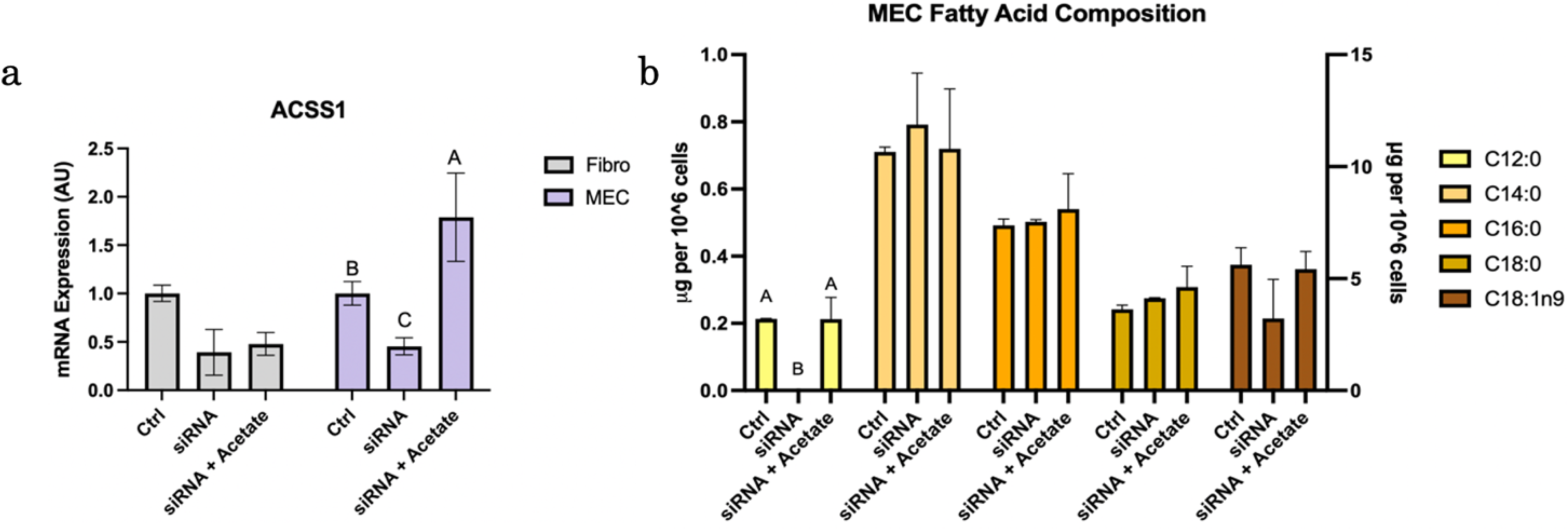
ACSS1 knockdown effect on gene expression and fatty acid profiling in MEC under control and acetate treatment conditions. (a) Relative ACSS1 mRNA expression in fibroblasts and MEC following transient esiRNA mediated knockdown of ACSS1 and supplementation with 40 mM of acetate (b) fatty acid composition of MEC following the knockdown of ACSS1 and the subsequent supplementation with 40 mM acetate. Significant differences (*p ≤ 0.05*) are indicated by different letters.

ACSS1 expression levels influenced lipid composition, specifically the medium-chain fatty acid C12:0, that was completely depleted following ACSS1 knockdown. However, acetate supplementation restored C12:0 levels despite the knockdown (**Figure 5b**), consistent with the ACSS1 expression pattern.

### 3.4. Metabolomic Analysis of MEC response to acetate

The general metabolic response of MEC to acetate was further investigated using LC-MS based metabolomics analysis. A total of 296 metabolites were identified (**Supplementary Table T2**), with nearly 70% of metabolites significantly altered. Of these, 105 metabolites increased in abundance following acetate treatment, with 67 showing statistically significant elevation. 189 metabolites decreased in response to acetate, with 134 significantly reduced (**Figure 6a**). Notably, the metabolites 3-nitro-L-tyrosine and gamma-L-glutamyl-L-alanine were detected exclusively in the acetate treatment group.

**Figure 6.**
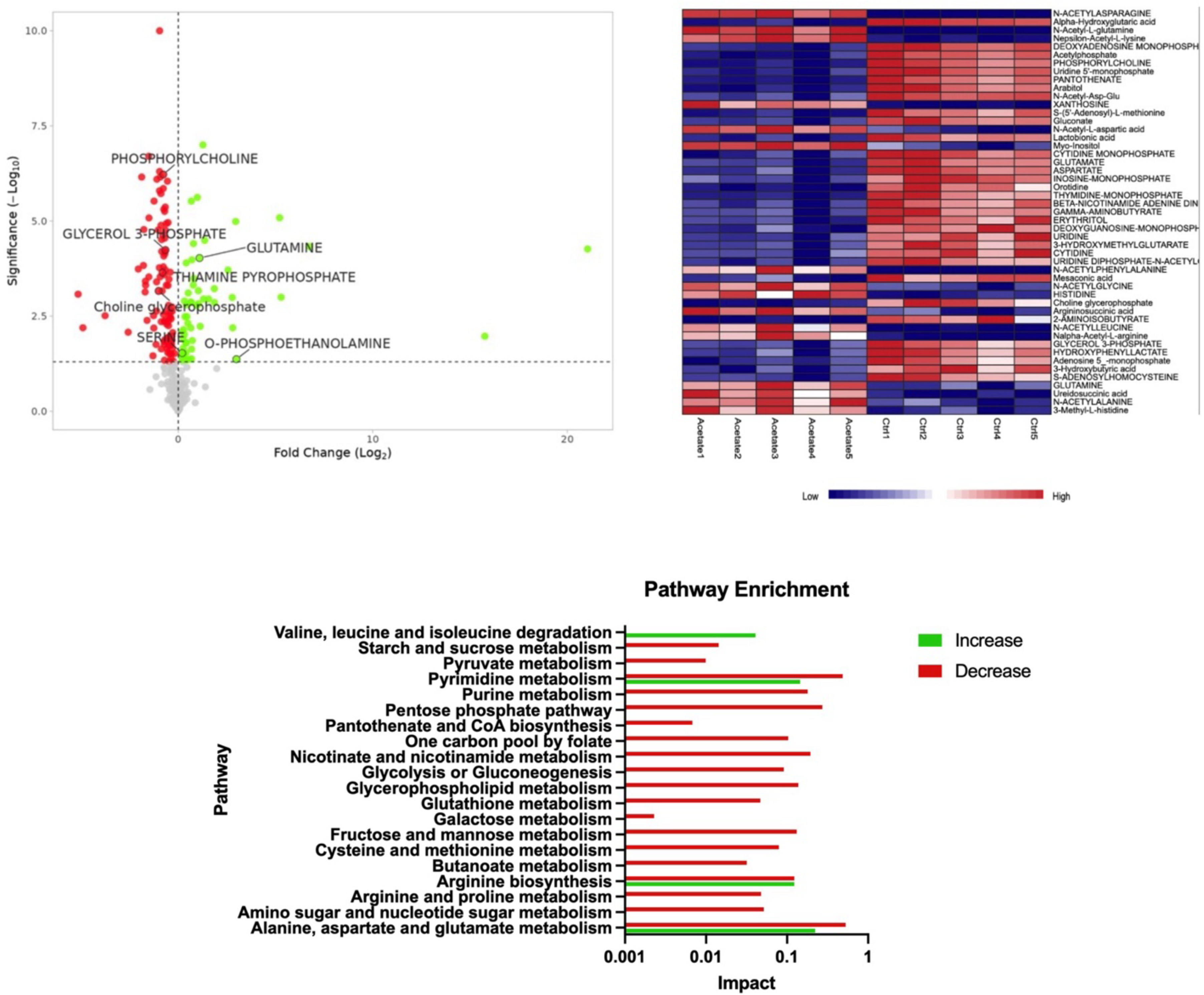
Metabolite abundance profiling and pathway enrichment analysis in MEC under control and acetate treatment conditions. (a) Volcano plot depicting the fold change (Log₂) versus significance (-Log₁₀ p-value) of all detected metabolites in acetate treated compared with control MEC. Red and green points represent significantly decreased and increased metabolites *(p ≤ 0.05)* respectively, while grey points indicate non-significant changes. Select metabolites of interest are labelled. (b) Upper 30% of metabolites that were significantly altered between treatments in MEC treated with and without 40 mM of acetate. Each row represents a metabolite, and each column represents a biological replicate. Colours indicate relative metabolite abundance (blue = low, red = high). (c) Pathway enrichment analysis performed using MetaboAnalyst 6.0, based on metabolites significantly altered by acetate treatment (*p ≤ 0.05*). Bars represent pathways enriched for increased (green) or decreased (red) metabolites. The x-axis shows pathway impact on a logarithmic scale.

To further investigate acetate-specific effects, we generated a heat map of metabolites. This visualization showed a clear separation between control and acetate-treated groups (**Figure 6b**). The full heatmap can be found in **Supplementary Figure S1**.

Pathway enrichment analysis revealed significant changes in multiple metabolic pathways following treatment (**Figure 6c**). Impact scores for each pathway were determined using topological analysis, with the direction of change based on the predominant pattern observed among metabolites in that pathway. Several amino acid metabolism pathways, including phenylalanine metabolism and alanine, aspartate and glutamate metabolism, showed significant alterations with different directional changes.

To further characterize the effect of acetate on MEC metabolism, we examined individual metabolite levels in MEC. By combining pathway enrichment analysis with the metabolites that contributed the most to treatment clustering (**Figure 6b**) we identified the pathways which can contribute or utilize intermediates in lipid metabolic pathways. These include the one carbon metabolism, TCA cycle, amino acid metabolism, glycerophospholipid metabolism, glycolysis/gluconeogenesis and the pentose phosphate pathway. Of those, the significantly different metabolites are presented (**Figure 7**). Thirteen free amino acids showed elevated levels in the acetate group compared with control, including alanine, arginine, isoleucine, glutamine, leucine, lysine, methionine, phenylalanine and serine, tryptophan, tyrosine and valine. Proline was the only amino acid that decreased in the acetate treated group (**Figure 7a**). In the glycerophospholipid metabolism pathway, acetate treatment resulted in increased levels of O-phosphoethanolamine and dihydroxyacetone phosphate, and decreased levels of glycerol 3-phosphate, citicoline, phosphorylcholine, and choline glycerophosphate (**Figure 7b**). Within the TCA cycle, levels of alpha-ketoglutarate, succinate, pyruvate, phosphoenolpyruvate, oxaloacetate, and thiamine pyrophosphate were all decreased in acetate treated cells relative to control (**Figure 7c**). In the one-carbon metabolism pathway, methionine and serine were elevated in the acetate group, while adenosine, (5’-adenosyl)-L-methionine (SAM) and S-adenosylhomocysteine were reduced (**Figure 7d**). Finally, in the pentose phosphate pathway, acetate treatment led to a decrease in 5-keto-D-gluconic acids while ribose, glyceraldehyde-3-phosphate, Sedoheptulose7-phosphate and 6- Phosphogluconate were increased (**Figure 7e**).

**Figure 7.**
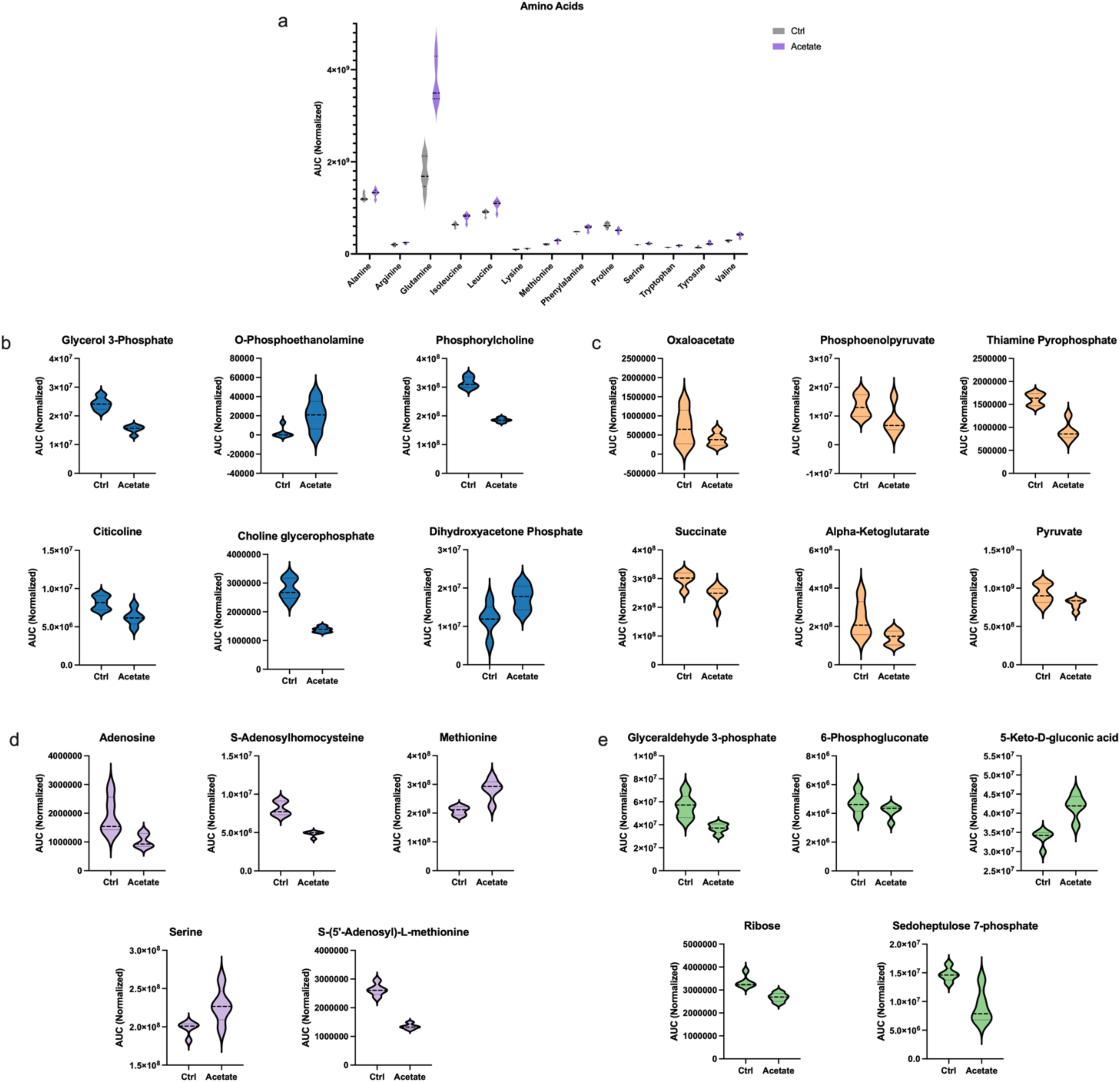
Distribution of metabolite concentrations in MEC under control and acetate treated conditions. Each plot reflects the relative abundance of specific metabolites measured by LC-MS, normalized to internal standards and total peaks, grouped by metabolic pathway: (a) Amino acids (b) Glycerophospholipid metabolism related metabolites (c) TCA cycle intermediates (d) One carbon metabolism metabolites (e) Pentose phosphate pathway metabolites. Violin plots depict the distribution of replicate values for each treatment. The central horizontal line indicates the median; shape width represents data density. All comparisons between control and acetate-treated conditions showed statistically significant differences (*p* ≤ *0.05*).

## 4. Discussion

Ruminant milk is highly enriched with short and medium chain fatty acids, a composition not found in milk of other mammalian species, or in other lipogenic tissues in ruminants such as the liver and adipose tissues ^4,29^. Although studies have been conducted over the years to reveal the mechanism underlying this phenomenon, it is still not clear how significant amounts of short and medium chain fatty acids are produced, accumulated and packed as triglycerides by bovine MEC and not by other fat producing bovine cells.

In the present study, we utilized a comparative approach to shed light on the pathways responsible for the different fatty acid composition of milk fat compared to other tissues. We used MEC lines (i.e. MacT and BME-UV1) from the bovine mammary gland and compared their basal and metabolically induced responses with that of fibroblasts isolated from the mesenchyme of lactating cow mammary gland. By directly comparing mammary epithelial cells and fibroblasts- both capable of triglyceride accumulation ^30^ - our approach enables the identification of mammary-specific regulatory mechanisms contributing to the unique composition of dairy fat. We expected that the fibroblast cell line will demonstrate adipose like properties in terms of lipogenic gene expression since adipocytes and fibroblast are capable of lipid synthesis and accumulation and share the same lineage ^31^. This comparative strategy provides a powerful framework for dissecting the pathways underlying the unique fatty acid composition of milk fat.

In a recent mega-analysis study, we compared gene expression patterns between mammary gland and adipose tissues and identified several expression differences that may affect the bovine mammary gland’s ability to synthesize short and medium chain fatty acids ^12^. For example, the ACSS1 gene, which converts acetate into acetyl-CoA, has a greater expression in the mammary gland compared to adipose tissue. Additionally, AGPAT1 and AGPAT6 genes are considered to be the dominant isoforms of the AGPAT family in the mammary gland ^21,12^. Notably, in the current study, basal gene expression of these genes was inconsistent with the expected *in vivo* patterns. Herein, bovine MEC showed only moderate ACSS1 expression, lower by two orders of magnitude from that of fibroblasts, and no notable enrichment of AGPAT1 or AGPAT6 compared to other isoforms. In accordance, fatty acid profile in MEC under basal conditions was different from that found in milk. Nonetheless, bovine MEC and fibroblasts lines exhibited significant differences from one another, both in terms of fatty acid composition as well as lipogenic genes expression. Moreover, ACSL1 and ACSL4, genes involved in the activation of long-chain fatty acids ^32^, were more highly expressed in fibroblasts under basal conditions, suggesting higher utilization of long chain fatty acids for triglyceride synthesis in accordance with the fatty acid composition found in extra mammary tissues ^4^.

We further studied the response of MEC to lipogenic building blocks and compared it to that of fibroblasts. In particular, acetate is one of the major lipogenic metabolites in ruminants, and it is found in relatively high amounts in the circulation due to forages fermentation in the rumen ^33^. It was hypothesized that the metabolic and molecular response to this common metabolite may differ between cell lines and may explain the differences in composition found *in vivo* between the mammary gland and other tissues. Therefore, we exposed MEC and fibroblasts to palmitate, acetate, or a combination of both. A consistent finding across both cell types used in the present study was the downregulation of FASN and ACC, two central enzymes of *de novo* fatty acid synthesis, in response to palmitate. This aligns with previous studies in MEC showing that palmitate, the end product of FASN, exerts negative feedback on its own synthesis by inhibiting FASN and ACC expression ^34^. Our results are also in accordance with previous studies in mammary cells demonstrating elevated FASN and ACC expression in response to acetate ^34,35^. However, beyond this shared response, the two cell types diverged in several aspects. Specifically, hierarchical clustering of gene expression responses (**Figure 2a**) revealed a clear separation between fibroblasts and MEC, underscoring the cell specific sensitivity to these lipogenic metabolites. Notably, only in MEC, palmitate induced ACSL1 expression, a response expected with the increased availability of long chain fatty acids. When acetate was supplemented to the media, even with supplementation of palmitate, ACSL1 expression in MEC was significantly and considerably decreased. Moreover, in MEC, acetate supplementation led to a significant upregulation of ACSS1, FASN, and DGAT, a response not observed in fibroblasts. These results indicate that MEC is highly sensitive to the presence of acetate, which is preferentially activated by ACSS1 and utilized for *de novo* short-chain fatty acid synthesis. At the same time acetate decreases the ability of the cells to utilize the supplemented palmitate as a substrate for triglyceride production by downregulating ACSL1. These results suggest that under physiological conditions, acetate, which is an abundant and highly available metabolite in ruminants, shifts the balance towards more effective utilization of short chain fatty acids for triglyceride synthesis, facilitating the production of the unique composition of diary fat. On the other hand, fibroblasts exhibited a downregulation of ACSS1 in response to palmitate, shifting the balance towards the activation of long chain fatty acids for triglyceride synthesis. These results also suggest that with elevated availability of lipogenic building blocks, MEC tend to activate more short chain fatty acids, and increase the capacity to incorporate them into sn-3 position on the triglyceride by DGAT. Indeed, in bovine milk, this position is occupied by short and medium chain fatty acids ^36^. Moreover, *in vivo*, the higher availability of acetate due to increased feed intake in dairy cows is associated with a shift in fatty acid composition of milk, increasing the proportion of short versus long chain fatty acids ^33^. Hence, our results suggest a molecular and metabolic mechanism which facilitates this unique feature of bovine milk, which is responsible for many of the organoleptic properties of bovine milk ^12^.

The specificity of the response of MEC to the acetate treatment is further supported by fatty acid composition data, which showed an acetate-dependent increase in medium chain fatty acid content (C10:0, C12:0, C14:0) in MEC, but not in fibroblasts (**Figure 3**). Considering that ACSS1 levels were higher in MEC receiving acetate treatment (Figure 2b) it is expected to produce more acetyl CoA, increasing the balance of intracellular acetyl-CoA and malonyl-CoA. A high acetyl-CoA to malonyl-CoA ratio has been shown to favour production of short and medium chain fatty acids ^22,37^. The significance of ACSS1 in this process was further demonstrated by a transient knockdown of this gene, resulting in near-complete loss of C12:0 in MEC, which was restored by acetate treatment alongside the recovery of ACSS1 expression (**Figure 5**). ACSS1 replenishment following acetate supplementation was an effect unique to MEC and was not found in fibroblasts. In addition, acetate treatment in MEC, but not in fibroblasts, resulted in an increase of triglyceride content (**Figure 4**), highlighting MEC elevated capacity to translate acetate availability into lipid synthesis. However, no change in total fatty acid composition was observed after palmitate supplement (**Figure 3**). This may suggest that in MEC, palmitate is directed to further metabolism pathways like elongation and desaturation ^26^, which can help conserving the composition of the fat. This could be attributed to the fact that fat produced by MEC has nutritional implications and therefore final lipid composition is maintained more strictly compared to other tissues.

To further understand the mechanisms induced by acetate in MEC we used metabolomics analysis to map the metabolic pathways modulated by acetate in this cell line. As demonstrated by a clear separation between control and acetate-treated MEC (**Figure 6b**) results indicate that acetate induced a broad metabolic shift beyond its expected role in promoting fatty acid synthesis and triglyceride accumulation. Notably, nearly 70% of detected metabolites were significantly altered by acetate (**Figure 6a**), underscoring the extensive metabolic changes triggered by acetate exposure in MEC. This distinction between treatments remains evident when focusing on specific metabolic pathways, including glycerophospholipid metabolism, glycolysis/gluconeogenesis, the TCA cycle, pentose phosphate pathway, and one-carbon metabolism (**Figure 6a**).

In our pathway analysis, one of the most prominent results is the effect of acetate on free amino acids. Thirteen amino acids were elevated following acetate treatment (**Figure 7a**), along with an increase in acetylated amino acid forms. This likely reflects a change in the flux of amino acids into the TCA cycle in accordance with the reduction of TCA cycle metabolites (**Figure 7c**) such as oxaloacetate, alpha-ketoglutarate and pyruvate. Moreover, elevated free amino acids under acetate treatment may support greater protein synthesis. This assumption is supported by previous work showing that acetate, especially in combination with leucine, enhances protein production *in vitro* in MEC ^38^ and *in vivo* by post rumen infusion of sodium acetate in lactating dairy cows ^39^. Although some studies suggested that acetate drives amino acid synthesis via the mTOR pathway ^35^, in the present study the mTOR pathway was not affected by acetate. Therefore, the results imply that acetate affects amino acid metabolically, however, further studied are warranted to confirm this hypothesis.

Higher free methionine levels in the cell and lower S-(5’-Adenosyl)-L-methionine (SAM), as reported herein (**Figure 7d**) may be a consequence of a change in phospholipid synthesis and degradation pathways (**Figure 7b**). More specifically, acetate treatment increased levels of o-phosphoethanolamine, an intermediate in phosphatidylethanolamine (PE) biosynthesis via the CDP-ethanolamine pathway, as well as decreased phosphorylcholine and citicoline, precursors in the phosphatidylcholine (PC) synthesis pathway ^40^. These findings suggest a shift in phospholipid composition, possibly favoring PE over PC. Since the conversion of PE to PC requires SAM as a methyl donor ^41^, favouring PE over PC production requires less methyl groups, sparing methionine and SAM utilization, explaining their increased concentrations. Moreover, a decrease of 3-glycerophosphate, a serine biosynthesis precursor ^42^, along with an increase in serine, further supports the shift toward PE synthesis. Higher PE content can induce fusion of intracellular lipid droplets and production of larger lipid droplets in the cells ^43,44,45^, a phenotype that was received in the present study following acetate treatment.

In summary, our comparative approach using MEC and fibroblasts provided valuable insights into the mammary-specific mechanisms of triglyceride synthesis and milk fatty acid composition. We observed that MEC, but not fibroblasts, responded to acetate supplementation with increased triglyceride accumulation and medium-chain fatty acid production, reflecting a distinct metabolic capability of MEC. These findings suggest that the mammary gland utilizes acetate differently from other tissues, potentially through higher expression and activity of ACSS1. Since acetate serves as a major metabolite in ruminant tissue metabolism, our findings demonstrate the bovine mammary gland’s enhanced capacity to utilize this lipogenic precursor through specific upregulation of lipogenic gene expression as well as broad metabolic reprogramming, ultimately contributing to the unique compositional characteristics of dairy fat.

## Acknowledgments

Nurit Argov-Argaman is the Baron de Hirsch Chair in Animal Husbandry.

This research was supported by The Israel Science Foundation (grant No. 2840/25)

## Abbreviations

ACC: acetyl-CoA carboxylase
ACLY: ATP citrate lyase
ACOX: peroxisomal acyl- coenzyme A oxidase
ACS: acyl-CoA synthetase
ACSL: acyl-CoA synthetase long chain
ACSS1: acyl- CoA synthetase short chain-1
AGPAT: acylglycerol phosphate acyltransferase
CK18: cytokeratin 18
DGAT: diacylglycerol acyltransferase
ER: endoplasmic reticulum
FACS: flow cytometry
FASN: fatty acid synthase
GC: gas chromatography
GPAM: glycerol-3-phosphate acyltransferase 1
GPAT: glycerol- 3-phosphate acyltransferase
MEC: mammary epithelial cells
MFG: milk fat globule
PBS: phosphate- buffered saline
PC: phosphatidylcholine
PCA: principal component analysis
PE: phosphatidylethanolamine
PEMT: phosphatidylethanolamine N-methyltransferase
PS: phosphatidylserine
PSD: phosphatidylserine decarboxylase
SAM: (5’-adenosyl)-L-methionine
TCA: tricarboxylic acid.

## CRediT author statement

Noam Tzirkel-Hancock: Data curation; Formal analysis; Methodology; Visualization; Writing - original draft; Writing - review & editing. Michal Rakover: Data curation. Roni Tadmor-Levi: Conceptualization; Funding acquisition; Data curation; Methodology; Writing - review & editing. Nurit Argov-Argaman: Conceptualization; Funding acquisition; Investigation; Methodology; Project administration; Supervision; Validation; Writing - review & editing

---

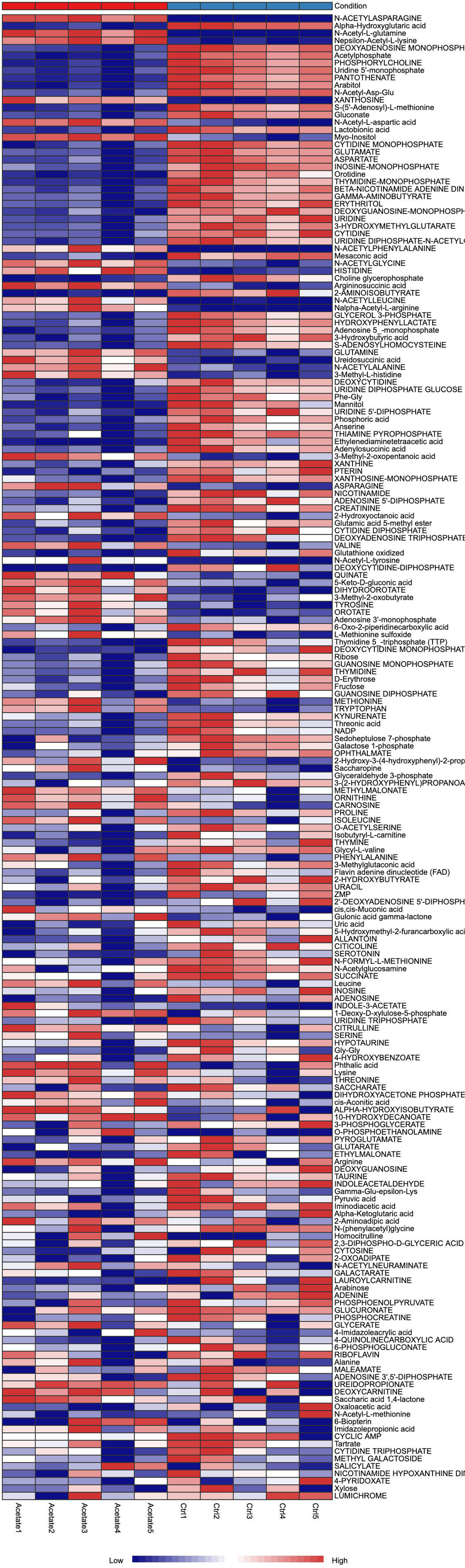

