## Supplementary Table T1 for "Lipogenic gene expression and substrate sensitivity in the bovine mammary gland shape milk fat composition"

**Supplementary Table T1.** List of Primers used for qRT-PCR

| **Gene Name** | **Accession Num.** | **Forward Sequence (5’-3’)** | **Reverse Sequence (5’-3’)** |
| --- | --- | --- | --- |
| 18S | NC_037354.1 | CGGCTACCACATCCAAGGAA | GGGCCCCGAAAGAGTCCTG |
| UXT | NM_001037471.2 | TGTGGCCCTTGGATATGGTT | GGTTGTCGCTGAGCTCTGTG |
| ACSS1 | NM_174746.2 | CCGATCAGGTCCTGGTAGTGA | CTCGGCCCATGACAATCTTC |
| DGAT | NM_174693.2 | CGACTCCTGGAGATGCTGTT | ATGCGGGAGTAGTCCATGTC |
| FASN | NM_001012669.1 | GCATCGCTGGCTACTCCTAC | GTGTAGGCCATCACGAAGGT |
| ACC | NM_174224.2 | AGCTGAATTTTCGCAGCAAT | GGTTTTCTCCCCAGGAAAAG |
| AGPAT1 | [NM_177518.1](https://www.ncbi.nlm.nih.gov/nuccore/NM_177518.1) | CTTCACAGCCCTACGTCGTT | AGCACCTCCATCATCCCAAG |
| AGPAT2 | NM_001080264.1 | GGCCTGATCATGTACCTGGG | CACATCGGTCATCACGCTCA |
| AGPAT6 | NM_001083669.1 | CGGAGTCTCCTTTGGTATCCG | CTCCATCCTCAAGGTAGCCC |
| ACSL1 | NM_001076085.1 | GGCCCATATGTTTGAGAGAGTT | GGGGAAGATGGTGGGTTGAA |
| ACSL4 | XM_015461583 | GCTGGGACCAAAGGATACGTAT | TCCAATCCTACAGCCATAGGTAAAG |
| GPAT3 | NM_001192514.3 | TCATCGTGCGCTACTGTGTT | TCCAACCAGGGTGGTTCCTA |
| ACOX1 | NM_001035289.3 | GTGGATATCAACAGCCCCGA | GAATCTGGAGGACTTTTTCCGT |
| ACOX2 | NM_001102015.2 | ACAGGTTCTCAGCACAGTCC | CTGCCTGGCATCCAAGAAGA |
| ACOX3 | NM_001103236.1 | CCCGAGGACTTGTTGGACTC | CTCGACTGAGCTTCTGGTGG |
| ACLY | NM_001037457 | CCTGCCATGCTCCAAGGAAA | GGGAGCAGACGTAGTCGAAG |
